# Decreased investigatory head scanning during exploration in learning-impaired, aged rats

**DOI:** 10.1101/2020.05.01.072249

**Authors:** Geeta Rao, Heekyung Lee, Michela Gallagher, James J. Knierim

## Abstract

“Head scanning” is an investigatory behavior that has been linked to spatial exploration and the one-trial formation or strengthening of place cells in the hippocampus. Previous studies have demonstrated that a subset of aged rats with normal spatial learning performance show head scanning rates during a novel, local-global cue-mismatch manipulation that are similar to those of young rats. However, these aged rats demonstrated different patterns of expression of neural activity markers in brain regions associated with spatial learning, perhaps suggesting neural mechanisms that compensate for age-related brain changes. These prior studies did not investigate the head scanning properties of aged rats that had spatial learning impairments. The present study analyzed head scanning behavior in young, aged-unimpaired, and aged-impaired Long Evans rats. Aged-impaired rats performed the head scan behavior at a lower rate than the young rats. These results suggest that decreased attention to spatial landmarks may be a contributing factor to the spatial learning deficits shown by the aged-impaired rats.

## Introduction

The effect of aging on spatial learning ability is variable in Long-Evans rats, as some aged rats (called aged-unimpaired, or AU) show no difference in performance relative to younger animals while other aged rats (called aged-impaired, or AI) show clear deficits in learning (Gallagher et al., 1993). Optimal execution of the classic Morris Water Maze task requires spatial mapping to locate the submerged platform relative to environmental cues (Hamilton et al., 2004; Knierim and Hamilton, 2011; Sutherland et al., 1987). If active attention to these cues is reduced in AI animals, this reduction may explain performance deficits in spatial tasks like the water maze. Defining epochs of attention to distal cues during the forward locomotion component of spatial exploration is difficult. However, certain investigatory behaviors such as head scanning (composed of lateral and vertical head movements) require suspension of locomotion followed by postural adjustments as the animal executes the behavior (Sinnamon et al., 1999), and thus these epochs are more detectable for analyses. Rearing, a behavior related to scanning, enables unique perspectives of the environment, particularly of distal cues, that are inaccessible during forward locomotion (Barth et al., 2018; Lever et al., 2006). Like rearing, head scanning provides unique views of the distal environment from distributed locations along the animal’s path as it pauses and alters head direction to gather information (Golani et al., 1993; Tchernichovski et al., 1998; Whishaw et al., 1994).

As head scanning has been associated with the new formation or strengthening of place fields in the hippocampal cognitive map (Monaco et al., 2014), alterations in this behavior could underlie aged-related impairments in spatial tasks. Prior studies have suggested that investigatory behaviors may be compromised in aged rats. Rearing in AI and AU rats is reduced in the plus maze and in the open field relative to young rats (Rosenthal et al., 1989; Rowe et al., 1998), but head scanning was not characterized in these studies. Orienting head movements in aged F344 rats have been described as sparse (Marini et al., 2006), although it was not reported if these animals showed learning impairments, and younger animals were not included for comparison. Increases in the back-and-forth head movements termed *vicarious trial and error* (Redish,2016; Tolman, 1939) have been reported in aged relative to young rats (Breton et al., 2015). However, this behavior occurs at decision points in relation to future fixed reward sites and thus may be qualitatively distinct from head scanning as observed during exploration. Recently, it was shown that AU rats showed head-scanning frequency comparable to young rats in a double-rotation, local-global cue mismatch manipulation that is amenable to identifying clear epochs of head scanning behavior (Branch et al., 2019; Haberman et al., 2019). This preservation of scanning behavior in the aged animals was correlated with increases in inhibitory interneuron activity as a potential compensatory mechanism to counteract the hyperactivity that has been reported in aged hippocampus (Branch et al., 2019). However, Branch et al. (2019) did not investigate head scanning behavior in AI rats. Reduced scanning behavior in AI rats may help to understand deficits in a wide range of spatial tasks, as it would suggest that reduced attention to spatial cues might be a contributing factor to these deficits. Alternatively, a finding of preserved scanning behavior in impaired animals might suggest that scanning in AI animals is not accompanied by compensatory neural mechanisms to adequately manage the influx of information. The current study was thus performed to investigate head scanning behavior in young (YG), aged unimpaired (AU), and AI (aged impaired) rats.

## Methods

### Subjects

Male Long–Evans rats (retired breeders) were obtained at 9 months of age from Charles River Laboratories (Raleigh, NC) and housed in a vivarium at Johns Hopkins University until behavioral assessment in the water maze at 22-26 months of age. Young rats obtained from the same source were housed in the same vivarium and tested at 4-6 months of age. All rats were individually housed at 25°C and maintained on a 12 h light/dark cycle. Food and water were provided ad libitum unless noted otherwise. The rats were examined for health and pathogen-free status throughout the experiments, and by necropsy after the experiment was finished. All animal care, housing, and surgical procedures were conducted in accordance with NIH guidelines using protocols approved by the Institutional Animal Care and Use Committee at Johns Hopkins University.

### Water maze testing

For the spatial version of the water maze task, the animals were given 3 trials per day for 8 days, in which the submerged escape platform was in a constant position in the tank (Branch et al., 2019; Haberman et al., 2019). The water temperature was 27° C. The release location in the tank was varied from trial to trial. If the rat did not locate the platform within 90 seconds, it was guided to the platform. All animals were allowed to remain on the platform for 20 seconds and then placed in a holding cage for 40 seconds, resulting in an intertrial interval of 60 seconds. Every sixth trial was a probe trial with the platform lowered to the bottom of the tank for the first 30 seconds, allowing assessment of search proximity. The platform was then mechanically raised to its normal, submerged position for the remainder of the trial. For analyses, a cumulative search error measure, reflecting the average distance of the rat from the platform, was obtained by averaging each block of 5 training trials. The learning index (LI), an average weighted proximity score during the probe trials (Gallagher et al., 1993), was used to classify the animals as impaired or unimpaired. On day nine, the animal was given six trials in which it had up to 30 seconds to locate the platform visible above the water level to screen for nonspecific task impairments such as swimming behavior and escape motivation. The location of the visible platform varied from trial to trial. Rats with health issues, such as pituitary tumors or signs of kidney impairment, were excluded from the study. Following water maze testing, the rats were placed on restricted food access to bring their body weight down to 85% while they were given foraging sessions (20 minutes per day for 10 days) in a cylindrical apparatus for chocolate pellets (Bio-Serv, Flemington, NJ). These foraging sessions provided pretraining to get the animals used to foraging for food reward before the subsequent training on the double rotation circular track.

### Double-rotation protocol

Animals were implanted with hyperdrives targeting the hippocampal CA3 subfield as previously described in detail for young rats (Lee et al., 2015). Neurophysiological data obtained from these animals will be presented elsewhere. The rats were trained in a double-rotation protocol (Knierim, 2002; Fig 1A). The double-rotation environment consisted of a wooden circular track (76 cm outer diameter, 56 cm inner diameter) with 4 distinctly textured quadrants: brown sandpaper, pebbled gray rubber mat, tan carpet pad, and gray duct tape with white labeling tape stripes. The circular track was mounted on a circular wooden platform at the center of a 2.7 m diameter curtained room. Global cues consisted of a cardboard circle with a white rectangular box on the floor below it; a blue surgical drape and white towel on an intravenous drip stand; a black and white striped card; a roll of brown wrapping paper; and a white card. The room was illuminated from above by a ring-light fixture in the center of the room. A white noise generator underneath the circular track masked directional sounds from outside the room. Over the course of 2-3 weeks, the rats’ weights were reduced further from the pretraining foraging sessions (described above) to 80-85% of free-feeding weights for YG and 75% for aged rats and they were trained to run clockwise on the circular track for chocolate pellets, with the local track cues and the global cues in a fixed configuration. Counterclockwise running was blocked with a piece of cardboard. When the rats were able to consistently complete 5 training sessions per day with at least 10 laps/session, the double-rotation protocol began. On each day of this protocol, three standard (STD) sessions with a cue configuration identical to training were interleaved with two mismatch (MIS) sessions. For the mismatch sessions, the global cues on the curtain were rotated clockwise and the local track cues were rotated counterclockwise by the same amount, representing a total cue mismatch of 45°, 90°, 135°, or 180°. The mismatch angles used on the first day in sessions 2 and 4 were randomly selected and the remaining two mismatch angles were used on the second day. An upper bound of 15 minutes on the track allowed for ample opportunity for exploration by animals that moved more slowly.

**Figure 1.**
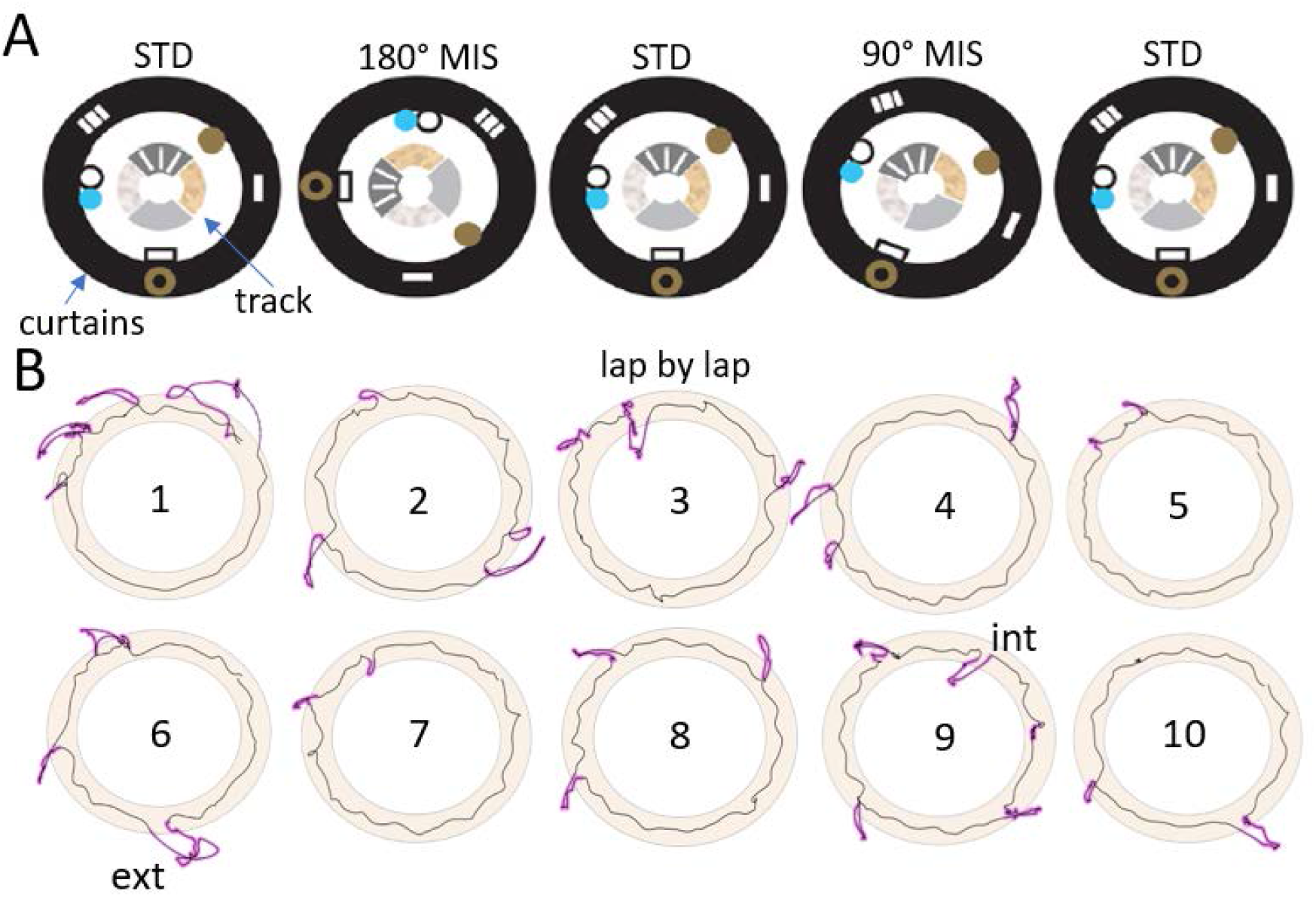
(A) The DR protocol showing the circular track with four distinctly textured quadrants within the curtained room. The daily protocol consisted of 3 standard sessions interleaved with 2 mismatch sessions. The example shows the standard cue configuration (std) in the first session followed by a mismatch session (mis) with a 90° clockwise rotation of the global cues, and an equivalent counterclockwise rotation of the track cues, resulting in a 180° mismatch. The second mismatch session shows the cue configurations corresponding to a 90° mismatch. Two of four possible mismatch angles (45°, 90°, 135°, and 180°) were randomly presented on day 1, followed by the remaining 2 angles on day 2 (not shown). (B) Enlarged images of the track alone, with black segments indicating the rat’s trajectories for 10 successive laps in a standard session. Quadrant textures are omitted. Purple segments are head scanning events. Examples of exteriorly directed (ext, lap 6) and interiorly directed (int, lap 9) head movements are labeled; only the more common exteriorly directed head scans were analyzed.

The animal’s position on the track (Fig 1B) was monitored at 30 Hz by overhead tracking of light emitting diodes attached to the rat’s head. Head scanning epochs were extracted from the rat’s trajectories during exploration by measuring lateral deviations from a running baseline distribution (see Monaco et al. [2014] for further details). Briefly, a running velocity threshold of 10° clockwise/s was used to detect forward movement around the circular track. Lateral head movements with a duration > 400 ms and a radial magnitude > 2.5 cm were classified as putative scanning events. Events that occurred within 400 ms of each other were merged and considered to be a single scan.

Algorithms to extract head scans from forward movement in the double rotation protocol were originally developed from a large database of young animals (Monaco et al., 2014), but the increasing dominance of other behaviors with aging complicated the analysis in the current study. For example, video inspection revealed that head movements directed toward the track interior typically involved grooming. Grooming behavior has a stereotyped structure, progressing cephalo-caudally from face-washing to a late phase involving lateral head movements as the animal grooms its trunk (Kalueff et al., 2016; Matell et al., 2006). Extended grooming was rare in young rats both in this study and previously (Monaco et al., 2014), but the aged animals in the current study groomed extensively. Aged rats have been shown to respond to novelty with increased grooming (Kametani et al., 1984; Rosenthal et al., 1989) and AI rats in particular may groom at the expense of investigatory behavior (Rowe et al., 1998), making the accurate isolation of investigatory head scanning bouts more difficult than in young animals. To focus on head scanning movements that are relatively free from grooming artifacts, we examined only exterior scans, i.e., head movements oriented laterally toward the curtained periphery where the distal cues were located. Interior head movements, directed toward the room center, were approximately one third as common as exterior scans and were more easily confounded with head movements associated with grooming, as the rats tended to face toward the interior of the track while grooming; thus, because of this ambiguity, these scans were excluded from analysis.

### Statistical Analysis

Data are reported as means and standard errors, except where stated otherwise for purely descriptive purposes. Two factor, mixed ANOVAs were performed on most of the data, with session number the within subject factor and age group the between subjects factor. Post hoc Tukey tests were used to compare specific differences between groups if there was a significant main effect or interaction revealed by the ANOVA. Effect size was calculated using Hedges’ *g* correction for sample size and unequal numbers per group. Correlations between learning index and scanning behavior were calculated using the Pearson product moment correlation.

## Results

### Water maze

Figure 2 shows the background water maze assessment of young and aged rats used in this study. Over the course of training, young rats were more proficient than aged rats in learning to locate the escape platform (Fig. 2A). A repeated measures ANOVA showed that the performance of both groups of rats improved over the course of training (F[3, 48] = 48.68, p = 0.001), but the young rats improved more rapidly as reflected by a significant interaction between age and trial block (F[3, 48] = 3.89, p = 0.014), and the young rats overall performed better than the aged rats (F[1, 16] = 5.41, p = 0.033). Figure 2B shows the learning index scores computed from a key measure of search accuracy during interpolated probe trials. Consistent with previous research in this model, the aged rats displayed a range of outcomes, with a subset of aged rats performing on par with young adults (aged unimpaired, AU) and a subset performing outside the range of young performance with a learning index score above 240 (aged impaired, AI). Cued training performance was not different among the three groups (mean ± se: young, 11.3 ± 2.4 s; AU, 8.7 ± 0.8; AI, 12.6 ± 1.7; [F(2,15) = 1.49, p = 0.256]), indicating that performance differences were confined to the spatial version of the water maze task.

**Figure 2.**
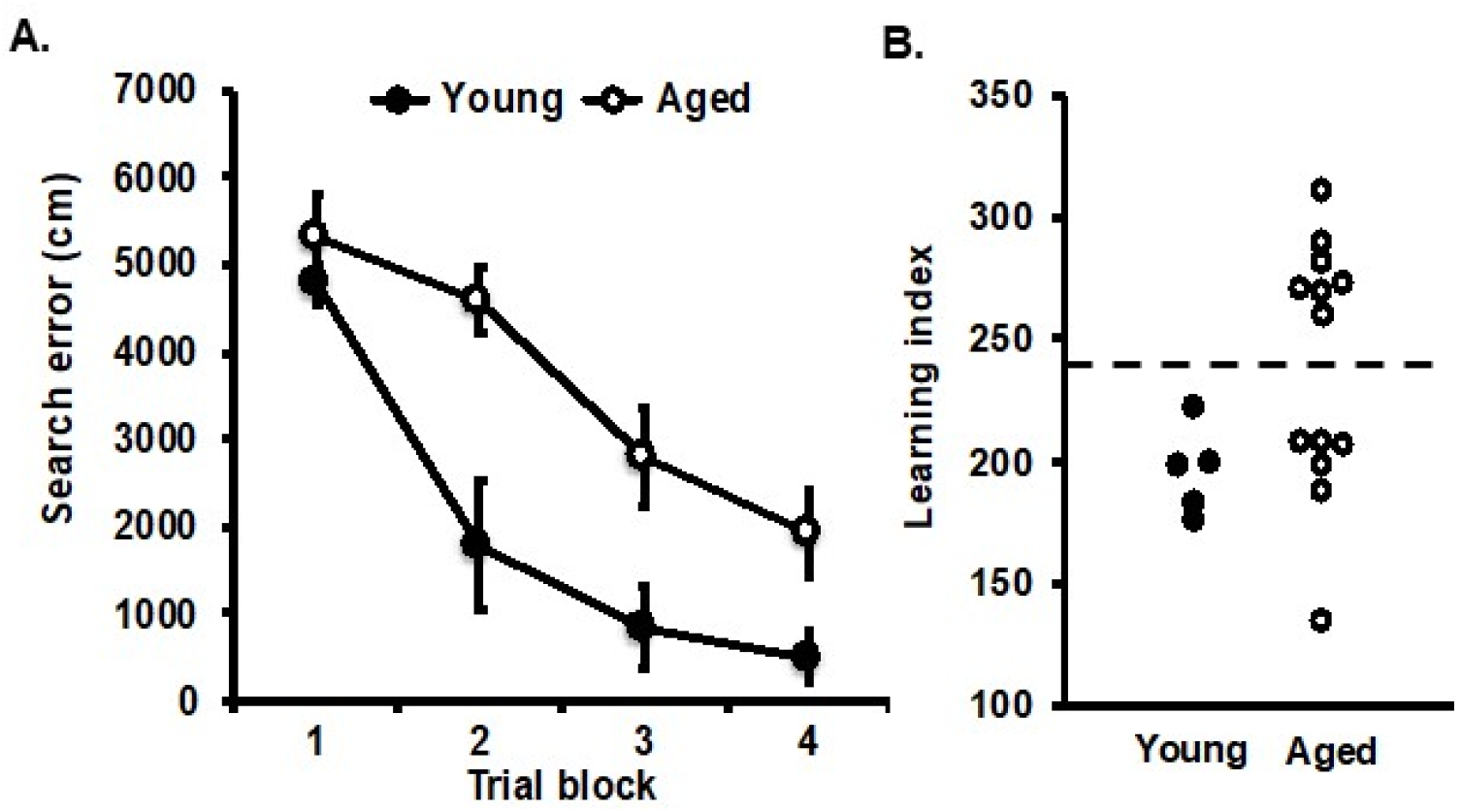
Background behavioral characterization of young and aged rats in the spatial version of the water maze. (A) Cumulative search error measure of learning during water maze training, which reflects the distance of the rat from the escape platform throughout its search, with higher numbers indicating worse performance. Each trial block consists of 5 training trials. (B) A learning index measure for each rat was derived from proximity of the rat’s search during probe trials interpolated throughout training, with lower scores indicating more accurate performance. Aged rats that performed on par with young adults were designated aged unimpaired (AU), and those that performed more poorly than young (learning index >240, dashed line) were designated aged impaired (AI).

### Double rotation

Behavioral data on the double rotation (DR) task were obtained from 5 YG, 6 AU, and 7 AI animals. Figure 3 shows examples from 3 animals from each age group (YG, AU, and AI) from a single standard session of the DR protocol. Variability in scanning behavior was present in all 3 groups, with some animals showing minimal scanning (left column) or substantial scanning (right column).

**Figure 3.**
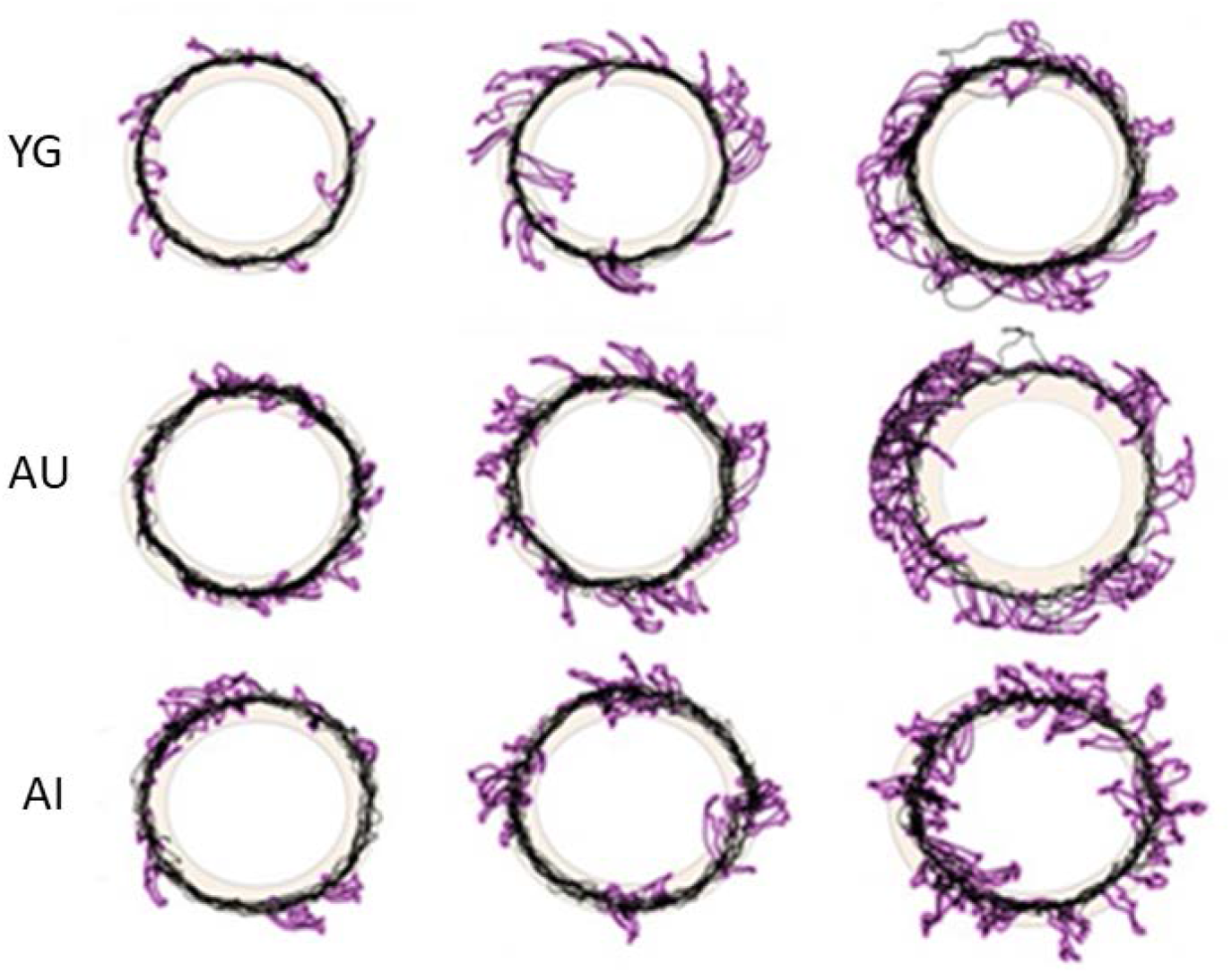
Variability in scanning behavior within age groups. Examples from 9 rats, 3 per age group, showing overlaid laps from a standard session on the circular track (quadrant textures not shown). Animals engaging in sparse (left column), intermediate (middle columns), or frequent (right column) lateral head movements were present in all age groups.

#### Session duration and time spent scanning

All animals from each age group completed 6-15 laps per session (mean ± s.d.,13.2 ± 1.7 laps) within the allotted time of 15 minutes, with almost all sessions including 9 or greater laps. There were no significant differences among age groups in the mean number of laps across the 5 sessions (Supplementary Table 1). However, AI rats took longer to complete the sessions than YG rats on Day 1 and they took longer than both YG and AU rats on Day 2 (Supplementary Table 1). To test for differences in baseline scanning behavior before the introduction of the novel, mismatch sessions, we analyzed data from the first session of Day 1 (the familiar, standard cue-configuration session). In this session, all 3 groups of rats spent approximately 25% of their time engaged in head scanning behavior (Supplementary Table 2). Thus, although the AI animals tended to circumnavigate the track more slowly, the opportunity to sample the environment from distinct locations over the course of the entire session was not different, as reflected by the similar mean number of laps and the similar proportions of time spent scanning among the age groups.

#### Scan magnitude and duration

We examined parameters of individual scans such as scan magnitude (the maximum radial distance from the averaged running trajectory along the track) and the duration of each individual scan. The magnitude of the head scans did not differ across age groups (Supplementary Figure 1; Supplementary Table 3). For scan duration, although there were no significant main effects of age or session, a significant interaction of age and session was found for Day 1 (Fig. 4, Table 1). Post hoc tests showed that the YG animals made significantly shorter duration scans than both the AI animals (sessions 4 and 5 of Day 1: session 4, p = 0.009, Hedges’ effect size *g* = 2.285; session 5, p = 0.0002, *g* = 3.191) and the AU rats (session 5 only; p = 0.0097, *g* = 2.504). On Day 2, there was a significant main effect of session, but no significant effect of age and no significant interaction. However, visual inspection of the data for Days 1 and 2 show similar trends, with both AU and AI rats differing little from YG rats in session 1 but steadily increasing their scan durations in session 2-4, whereas YG rats maintained a mostly constant duration. It is unclear if the longer duration scans reflect increased inspection of cues, slower head movements related to fatigue over the course of the experiment, or a combination of factors in the aged animals.

**Table 1.** Scanning duration.

| Table 1. Scanning duration |  |  |  |  |  |  |
| --- | --- | --- | --- | --- | --- | --- |
| DAY 1 |  |  |  |  |  |  |
| Source |  | SS | df | ms | F | p |
|  | AGE | 52.496 | 2 | 26.248 | 2.802 | 0.093 |
|  | ERROR<sub>b</sub> | 140.531 | 15 | 9.369 |  |  |
|  | SESSION | 2.448 | 4 | 0.612 | 0.887 | 0.476 |
|  | AGE x SESSION | 13.847 | 8 | 1.731 | 2.509 | 0.018 |
|  | ERROR<sub>w</sub> | 41.396 | 60 | 0.690 |  |  |
| DAY 2 |  |  |  |  |  |  |
| Source |  | SS | df | ms | F | p |
|  | AGE | 31.726 | 2 | 15.863 | 2.226 | 0.142 |
|  | ERROR<sub>b</sub> | 106.907 | 15 | 7.127 |  |  |
|  | SESSION | 8.922 | 4 | 2.230 | 2.968 | 0.025 |
|  | AGE x SESSION | 9.869 | 8 | 1.234 | 1.642 | 0.127 |
|  | ERROR<sub>w</sub> | 45.09 | 60 | 0.751 |  |  |

**Figure 4.**
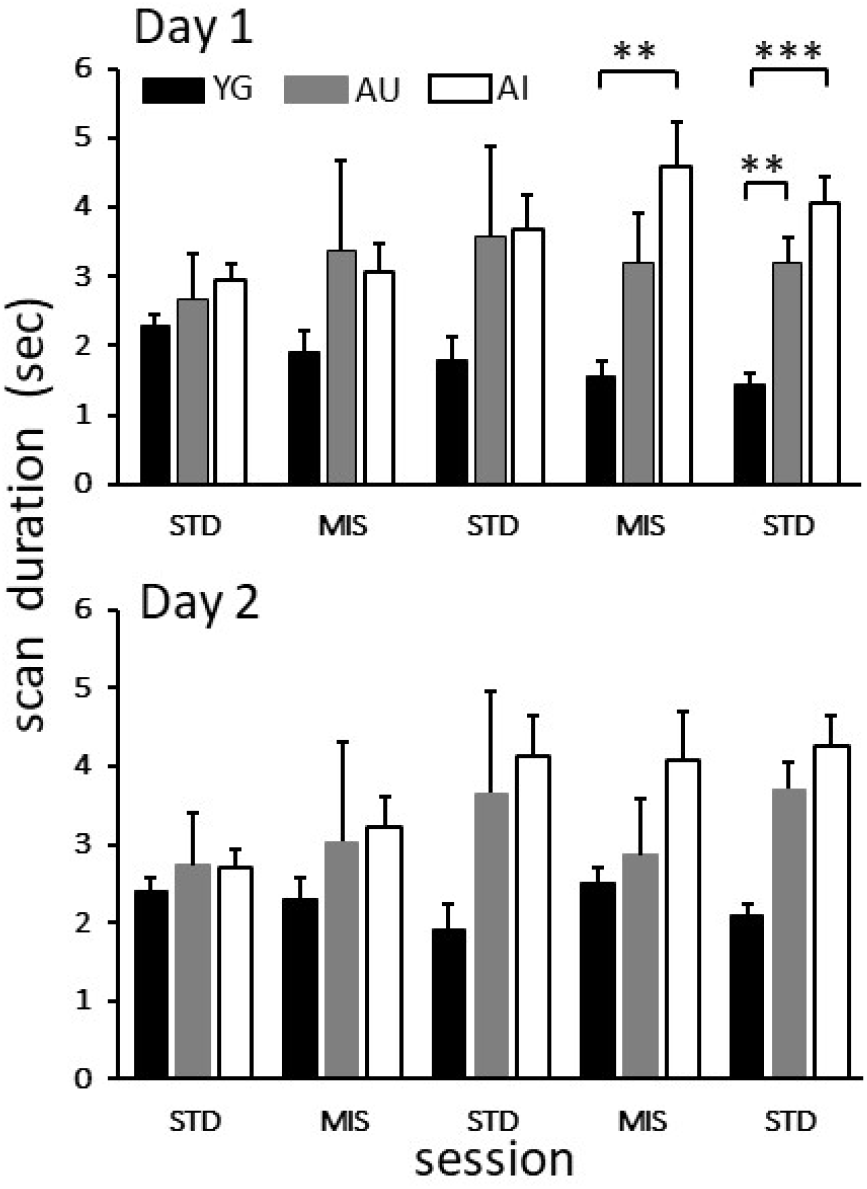
Scan duration across the 5 sessions for Day 1 (top) and Day 2 (bottom). No main effects were significant, but there was a significant interaction between age and session. Post hoc comparisons (Tukey HSD) indicated that in session 4, YG rats made significantly shorter duration scans than the AI but were not different from the AU rats (mean ± se: YG: 1.562 ± 0.205; AU: 3.198 ± 0.720; AI: 4.588 ± 0.205). In session 5, scans in the YG rats were shorter than those in the AU and AI animals (mean ± se: YG: 1.456 ± 0.153; AU: 3.191 ± 0.358; AI: 4.044 ± 0.381). Similar statistics for Day 2 did not reveal any significant differences across age groups for scan duration. An overall effect of session was found, with longer duration scans occurring in the later sessions. No significant interaction of age group and session was evident. Note that the large error bars for AU rats in sessions 2 and 3 result from a single animal that spent a large amount of time with its head off the track during a scan. ** p < 0.01, *** p < 0.001 post hoc Tukey tests. Wide brackets refer to comparison between YG and AI groups. Narrow bracket refers to YG and AU comparison.

#### Scanning rate

Although the proportion of time performing scanning within a session was roughly equal among age groups, there were significant age group differences in the frequency of head scanning. Scanning rate (the number of scans per minute) was analyzed for each of the 5 daily sessions on both days (Fig. 5). For Day 1, there were significant main effects of age group and session, with an interaction that just missed statistical significance (Table 2). Post hoc Tukey tests showed that there were no significant differences among the 3 age groups in the first standard session (i.e., the familiar cue configuration prior to any cue manipulations). However, for the remaining sessions, including the first mismatch session ever experienced by the rats (Session 2), the YG rats scanned at a significantly higher rate than the AI rats (Fig. 5). This result suggests that scanning behavior, a means of sampling environmental information, is attenuated in the AI rats.

**Table 2.**
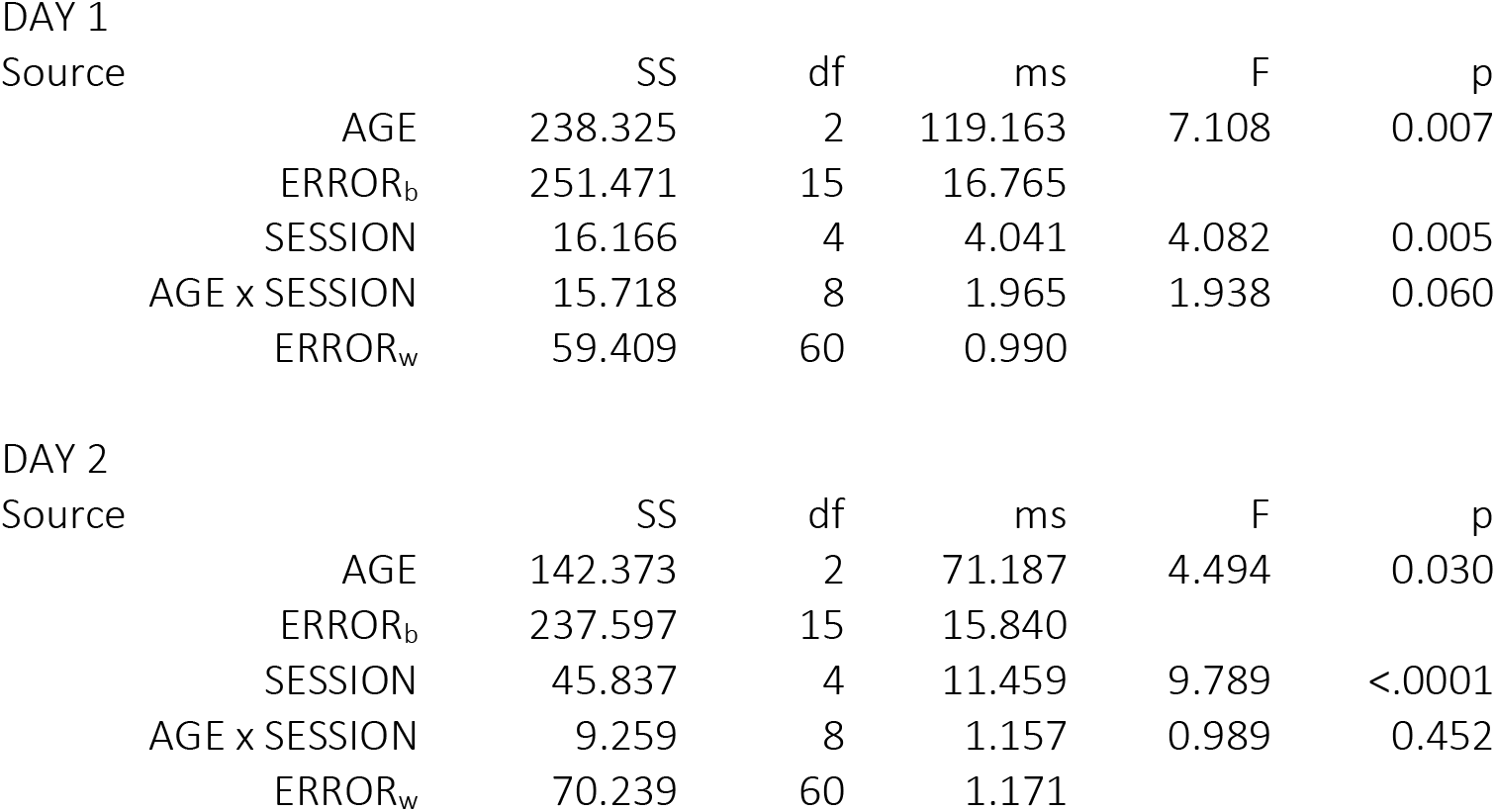
Scanning rate.

**Figure 5.**
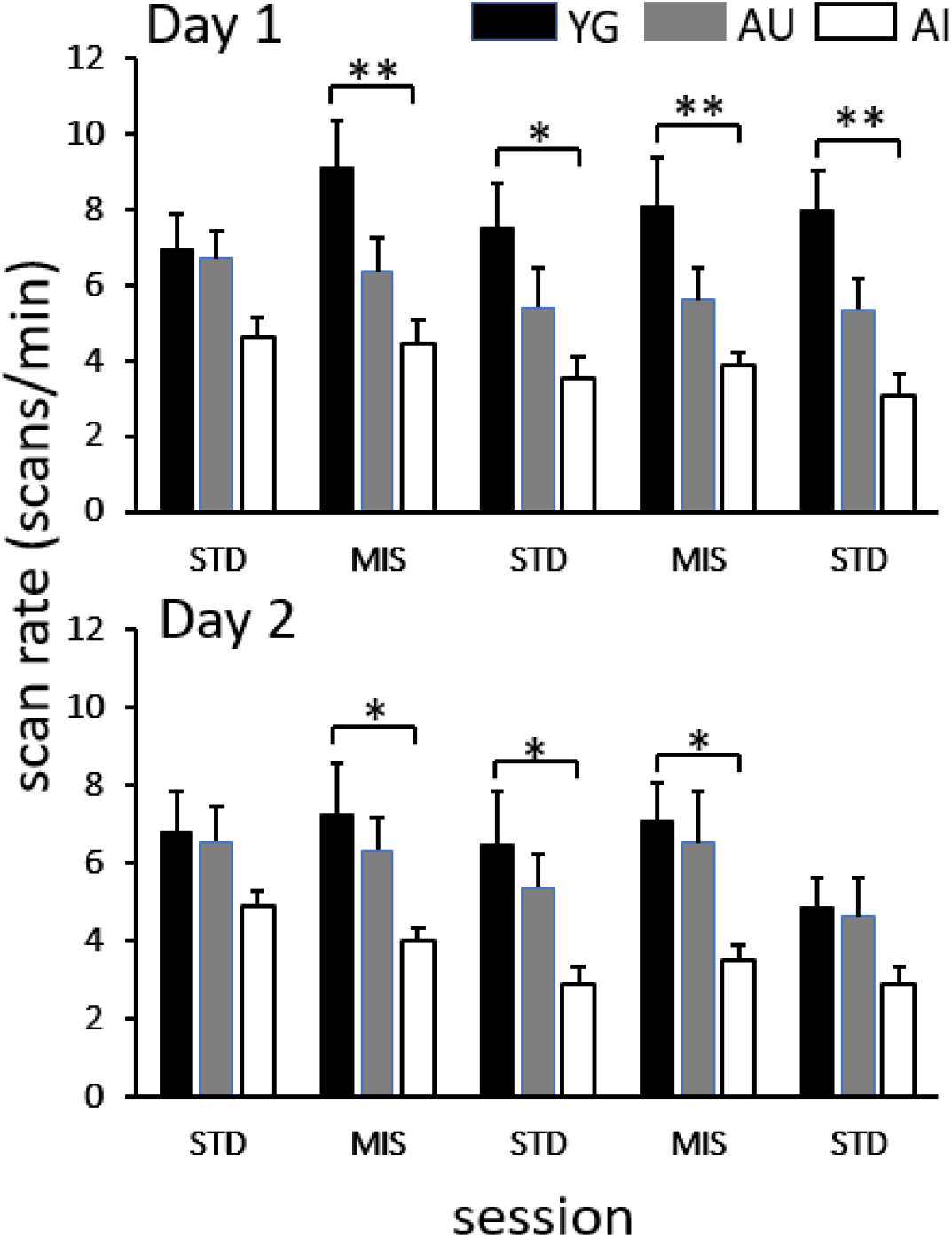
Scanning rates (scans/min) across 5 sessions of the DR protocol. Three standard (STD) sessions interleaved with two mismatch (MIS) sessions are shown for Day 1 (top) and Day 2 (bottom). On Day 1, YG rats showed significantly elevated scanning rates relative to the AI rats in sessions 2-5. On Day 2, YG rats scanned significantly more than the AI group on sessions 2, 3, and 4. No significant differences in scanning rate between the YG and AU rats were found either on Day 1 or Day 2. YG: Black, AU: Gray, AI: White. * p < 0.05, ** p < 0.01, post hoc Tukey tests. Hedges’ *g* effect sizes for YG vs. AI, Day1: session 2, 2.100; session 3, 1.936, session 4, 2.125, session 5, 2.682. Day 2: session 2, 1.672; session 3, 1.694; session 4, 2.277. Brackets refer to comparisons between the YG and AI groups.

One possibility is that the YG rats scanned more than the AI rats in the first mismatch session because they were initially presented, by chance, with a larger average cue mismatch than the AI group. On the contrary, the YG experienced less of a cue mismatch than AI rats in Session 2 (Supplementary Table 4). Hence, the significantly elevated scanning rate in YG relative to the AI group was not due to a larger mismatch angle. (The mean angles presented for other sessions during Day 1 and Day 2 were not different across age group; Supplementary Table 4).

The reduction in scanning rate in the AI rats was apparent also on Day 2. An ANOVA demonstrated significant main effects of age group and session; however, unlike Day 1, on Day 2 the interaction between age and session was not close to significance (Table 2). Post hoc tests showed that YG rats scanned significantly more than the AI group on sessions 2, 3, and 4 (YG vs AI: session 2, p = 0.036; session 3, p = 0.028, session 4, p = 0.045). In contrast, no significant differences in scanning rate between the YG and AU rats were found either on Day 1 or Day 2. Reduced AI scanning rates are consistent with the hypothesis of deficits in cue-sampling investigatory behavior in these animals.

### Correlation between scanning behavior and water maze performance

We examined whether the performance by aged animals in the spatial and cued versions of the water maze task was correlated with subsequent scanning behavior on the DR track. A negative correlation between the learning index and scan rate in the DR apparatus was found in the aged animals (Fig. 6A). That is, a smaller learning index, reflecting better performance in the spatial task, was significantly associated with higher scanning rates, a measure of attentive investigation in the DR protocol. Figure 6 shows that this negative correlation was true both for the first DR standard session on Day 1 (r= −0.708, p = 0.007) and for the largest (180°, Day 1 or Day 2) mismatch condition session that corresponded to the mismatch magnitude investigated by Branch et al. (2019) (r = −0.564, p = 0.045). (The 90° and 135° mismatch sessions in the present experiment also showed significant negative correlations, whereas the smallest mismatch angle of 45° showed no significant correlation; data not shown.) In contrast, for the cued version of the task (Fig. 6B), there was no correlation between the average latency to find the visible platform and subsequent scanning behavior (1^st^ std session: r = 0.071, p = 0.818; 180° mis: r = 0.050, p= 0.871). Unlike the aged group, YG animals did not show significant correlations between scanning rate and learning index in the spatial task (std: r = 0.070, p = 0.911; mis: r = 0.061, p = 0.923). However, a significant positive correlation was found between YG scanning rates in the standard session and latency to visible platform (r = 0.891, p = 0.041, data not shown), but this was not evident for the mismatch session (r = 0.729, p = 0.162).

**Figure 6.**
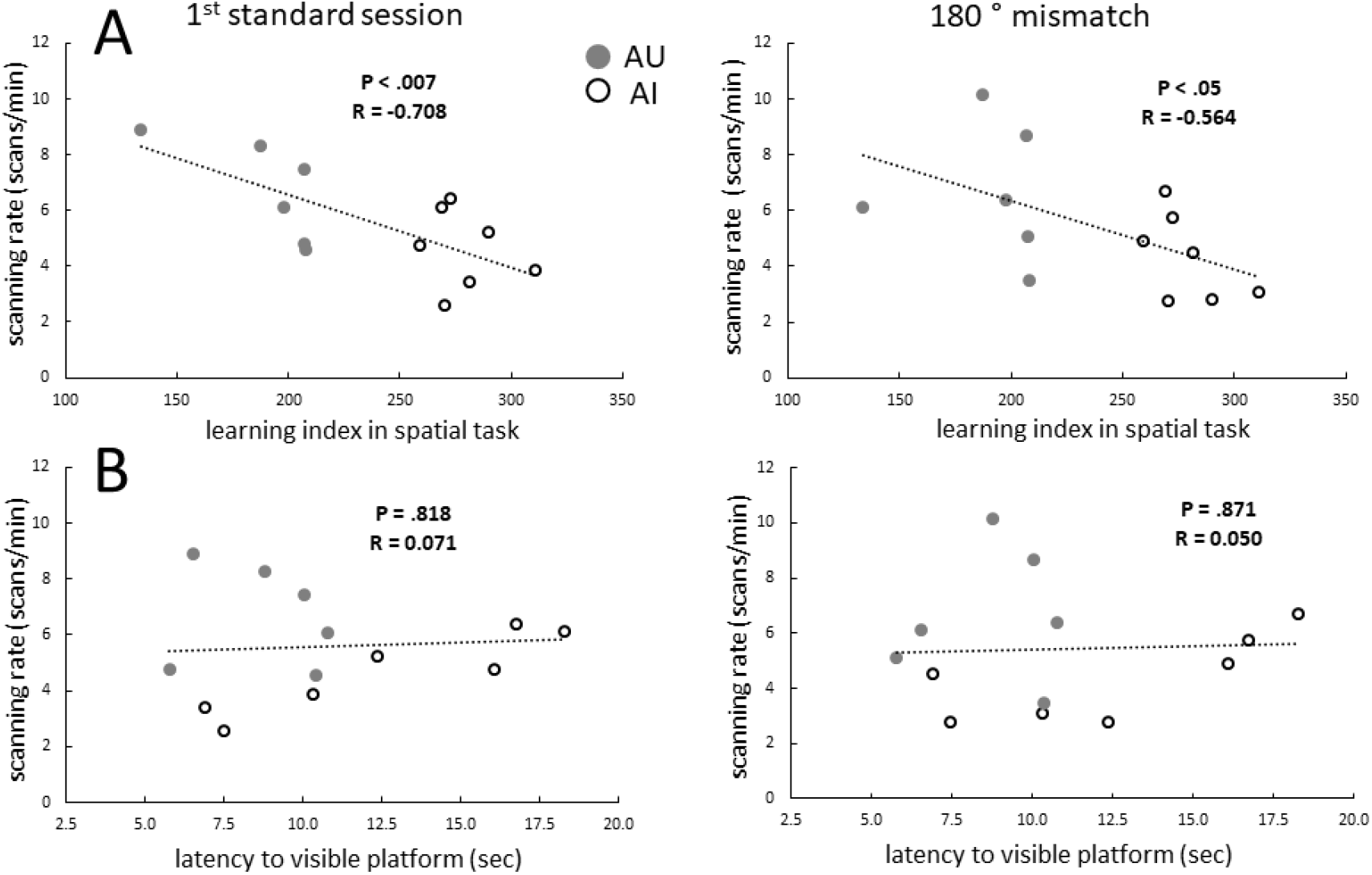
Relationship between aged animals’ water maze performance and scanning behavior in the DR protocol. (A) Correlations between the learning index from the spatial version of the water maze and scanning rate for the first DR standard session from Day 1 (left) and the 180° mismatch session (right). Both linear regressions were significant. (B) Correlations of the average latency in the visible platform task and subsequent DR scanning rate were not significant, either for the first standard session (left) or the 180° mismatch session (right). AU: Gray circles; AI: White circles.

## Discussion

Head scanning during exploration is a potential mechanism for rapidly binding information about the external world to a representation of the animal’s current location on an internal cognitive map (Monaco et al., 2014). A previous study investigating head scanning in aged animals (Branch et al., 2019) used a simple double-rotation protocol in which YG and AU rats were presented either a baseline session or a single, maximal (180°) cue mismatch session on the test day. In that study, both AU and YG animals showed similarly elevated scanning rates in the DR session compared to baseline. Furthermore, the presumed ability to detect environmental change, as reflected by scanning, was accompanied by an increase in zif268 activation in the hippocampal principal cells in both groups. However, an increase in hippocampal inhibitory interneuron function, measured by GAD1 activity, was selectively increased only in the AU animals, suggesting a potential compensatory mechanism to moderate age-related hyperactivity of hippocampal principal cells responding to environmental change (Branch et al., 2019). The purpose of the Branch et al. (2019) study was to compare YG rats with AU rats with similar learning performance, in order to look for such compensatory mechanisms in the AU brains. Accordingly, that study did not investigate the head scanning behavior of AI rats, and it was therefore unknown whether alterations in head scanning behavior may be a contributing factor to the learning impairments shown by AI rats. The present report is the first to compare head scanning behavior in learning-impaired, aged rats to that observed in unimpaired animals. We show that some parameters of this investigatory behavior appear to be significantly altered in AI rats compared to YG and AU rats.

Two measures of investigatory behavior, scan duration and scan rate, were different in the YG group compared to the AI animals, with intermediate values for AU animals that were usually not significantly different from the other groups. Scan duration (average seconds per scan) was significantly shorter in the young rats compared to the AI group, with the AU rats showing intermediate values. Age differences in this parameter are not straightforward to interpret, as scan duration measures the time from onset of the scan to a return to the baseline trajectory on the track, which may include epochs of immobility with the head laterally displaced. The second measure, scan rate, is depressed in the AI rats, consistent with a prior study indicating fewer head scans in aged rats (Marini et al., 2006), although that study did not address deficits in spatial learning. In the highly familiar condition prior to any cue manipulations, although a reduced scan rate was evident in the AI rats, it was not significantly different from the other groups. However, significant differences in scan rate emerged upon counter-rotations of the global and local cues, as the young rats elevated or tended to maintain their scan rates in the presence of cue alterations but the AI animals’ scan rates decreased across sessions. A similar pattern persisted on the second day until the last session. For this session, young animals were no longer scanning significantly more than the AI group, which may reflect habituation to the repeated cue re-arrangement. The AU scanning rates did not increase in the first mismatch condition of our study, whereas Branch et. al (2019) found an increase in scanning. However, the Branch study presented a single maximal mismatch angle of 180° to all animals. In contrast, in the present study, the presentation order of mismatch angles was randomized and, by chance, the first mismatch angles the AU animals experienced were substantially smaller than those of the other two age groups (Supplementary Table 4). However, even with this more subtle environmental change, the scan rate for the AU animals was intermediate between the YG and the AI animals, and was not significantly different from the YG animals.

Performance of all aged animals in the spatial version of the water maze was correlated with head scanning rate in the double rotation task. This result is consistent with reports that animals engage in cue-sampling behaviors similar to head scanning in the water maze task. The animal assesses the location of the submerged platform relative to distal cues during initial phases of the task (Hamilton et al., 2004; Knierim and Hamilton, 2011; Sutherland et al., 1982, 1987). Upon locating the hidden platform, a solid substrate like the circular DR track, rats engage in rearing and body turns and continue inspection of the distal environment (Sutherland and Dyck, 1984). Interestingly, behavior on the platform described as “orienting-like”, including head and body turns, appears to be altered in aged rats (Schulz et al., 2007). Analyses of water maze data, including the platform portion of the task, may reveal differences in how the AI animals survey their surroundings compared to AU and YG animals. However, we did not record video data from the water maze with sufficient resolution to determine behaviors that correspond precisely to the head scanning behavior on the circular track (which required the high-resolution tracking of head direction with colored LEDs on the rat’s head; Monaco et al., 2014), and we did not video-record any of the data on the escape platform. Thus, it remains for future investigations to determine what behaviors during the swimming and platform-occupancy phases of the water maze correspond to the head-scanning behaviors seen on the circular track, and whether these behaviors correlate with learning in the same way as head-scanning on the track. This would be difficult using LEDS in the water, but perhaps sophisticated algorithms like DeepLabCut (Mathis et al., 2018) might be utilized in the future for this purpose.

Reduced scanning in the AI animals is of interest in the context of exploration and information gathering. As the rat circumnavigates the circular track in our protocol, it samples the environment with unique perspectives only available during stops, as it cannot make head scanning movements while running (Sinnamon et al., 1999). The animal therefore may be incorporating distinct information about relationships between, or distances to, cues that is only accessible during scanning. For example, on the track, the rat cannot measure the distance to the global cues at the room periphery directly using path integration, but motion parallax via head movements during scanning may allow it to make an estimation (Legg and Lambert, 1990). One hypothesized function of intermittent locomotion, the interleaving of forward translation and pauses (Kramer and McLaughlin, 2001), is a moderation of the amount of incoming information during exploration (Benjamini et al., 2011). Interestingly, there appears to be an intrinsic upper limit on the number of stops an animal makes during exploratory excursions prior to returning to its home base (Golani et al., 1993). It has been further suggested that during pauses in exploration, a representation of place is stored in a buffer in the hippocampus and the upper bound represents the maximal number of these representations that can be stored upon the animal’s return to its home base (Jensen and Lisman, 1996). Impaired scanning in the aged-impaired rats may therefore compromise their ability to compartmentalize environmental information, particularly relating to distal cues, into manageable chunks of spatial information for subsequent storage and retrieval.

Head scanning may also be critical for the initial formation of cognitive maps. Animals and robots use path integration to create an initial, rudimentary map of location by integrating a movement vector (speed and direction) to update a representation of allocentric position as the animal makes small excursions from a home base in a novel environment (Thrun et al., 2005; Savelli and Knierim, 2019(Savelli and Knierim, 2019). During these excursions, through a mechanism similar to Hebbian learning, intermittent pauses and head scans may bind landmark information to specific locations on the map, thus allowing the internal map to remain stable relative to the external world. Ever-expanding excursions from the home base allow the animal to eventually create a global map of the larger environment. According to this scenario, spatial learning deficits in aged rats may in part arise from the altered head-scanning behavior, perhaps coupled with deficits in synaptic plasticity (Adams et al., 2001; Burke and Barnes, 2006; Foster, 2012) or neuronal excitability (Haberman et al., 2017; Koh et al., 2010; Wilson et al., 2005, 2003), thus impeding the formation of stable, reliable maps that are necessary for the proper execution of hippocampus-dependent learning tasks.

A prior finding that place field formation or strength is influenced by head scanning (Monaco et al., 2014) suggests that incoming environmental information during scanning behavior is being incorporated into a hippocampal representation of place. Other methods have been used to assess activity in hippocampal and related cortical regions related to investigatory behavior in a simplified version of the DR protocol (Branch et al., 2019; Haberman et al., 2019). In these studies, scanning rates in AU animals and YG animals were comparable in standard sessions, and were similarly elevated during the single 180° mismatch condition relative to baseline. This result indicated preserved recognition by the AU animals of spatial environmental change. The increase in scanning rate in the mismatch session was accompanied by similar increases in the activity of excitatory neurons in the hippocampus (Branch et al., 2019) and other cortical regions (Haberman et al., 2019) for both YG and AU rats, as measured by zif-268 expression. However, the expression of the inhibitory gene marker for the GABA-enzyme GAD1 was increased in the hippocampus in AU rats only (Branch et al., 2019). Because AI rats are known to have abnormally high levels of hippocampal activity (Wilson et al., 2003, 2004, 2005), which are correlated with poorer performance on the water maze (Haberman et al., 2017), Branch et al. (2019) suggested that the increase in interneuron activity in the mismatch sessions in the AU rats may be a compensatory mechanism to limit hippocampal neuronal hyperexcitability in the mismatch session. This compensation presumably keeps the population activity level of hippocampal neurons in AU rats in the normal range displayed by YG rats. The upregulation of inhibition in AU rats is functionally significant as disrupting it impairs spatial memory (Koh et al., 2020).

Although the current study is the only explicit characterization of head scanning behavior in AI animals, a prior finding in AI animals may nonetheless be informative. A number of interneuron changes are found across hippocampal subfields in aged vs. young animals, but a subpopulation of hilar interneurons is particularly affected in AI animals (Spiegel et al., 2013). The number of somatostatin-immunoreactive interneurons in the hilus of the dentate gyrus was selectively reduced in AI vs. YG and AU rats, and this reduction was correlated with spatial memory in the aged groups. The locus of differential change, specifically targeting a hilar subpopulation of interneurons in AI animals, is particularly intriguing in the context of investigatory behavior. The balance of excitation and inhibition in the dentate gyrus circuit is in part controlled by norepinephrine release in the hilus (Harley, 2007). Infusions of norepinephrine into the hilus promote rearing (Flicker and Geyer, 1982), a behavior closely tied to scanning, and “diversive” exploration, the sampling of a wide range of stimuli (Berlyne, 1960). Further study may reveal how the effect of a reduced influx of environmental information during scanning in AI rats may be compounded downstream by age-related changes in hippocampal circuitry to generate deficits in spatial memory.

## Acknowledgments

We thank Audrey Branch and Ming Teng Koh for comments on the manuscript and help in data analysis. Robert McMahan, Andrew Sherwood, Nicholas Lukish, Arjuna Tillekeratne and Kimberly Nnah assisted in running the behavioral experiments. Joseph D. Monaco developed the SCANR software for behavioral analyses. Chia-Hsuan Wang assisted in creating the aging database and subsequent analyses. Vyash Puliyadi provided further programming assistance. Supported by NIH grant P01 AG009973.

## Disclosure statement

Dr. Gallagher is the founder of AgeneBio Incorporated, a biotechnology company that is dedicated to discovery and development of therapies to treat cognitive impairment. She has a financial interest in the company and is an inventor on Johns Hopkins University’s intellectual property that is licensed to AgeneBio. Otherwise, Dr. Gallagher has had no consulting relationships with other public or private entities in the past 3 years and has no other financial holdings that could be perceived as constituting a potential conflict of interest. All conflicts of interest are managed by Johns Hopkins University. Ms. Rao, Dr. Knierim and Dr. Lee have no conflicts of interest to declare.

## CRediT authorship contribution statement

Geeta Rao: Conceptualization investigation, Formal analysis, Visualization, Writing - original draft. Heekyung Lee: Methodology, Investigation, Writing - review & editing Michela Gallagher: Conceptualization, Funding acquisition, Writing - review & editing. James Knierim: Conceptualization, Funding acquisition, Writing - original draft, Supervision.

## SUPPLEMENTARY MATERIAL

**Supplementary Figure 1.**
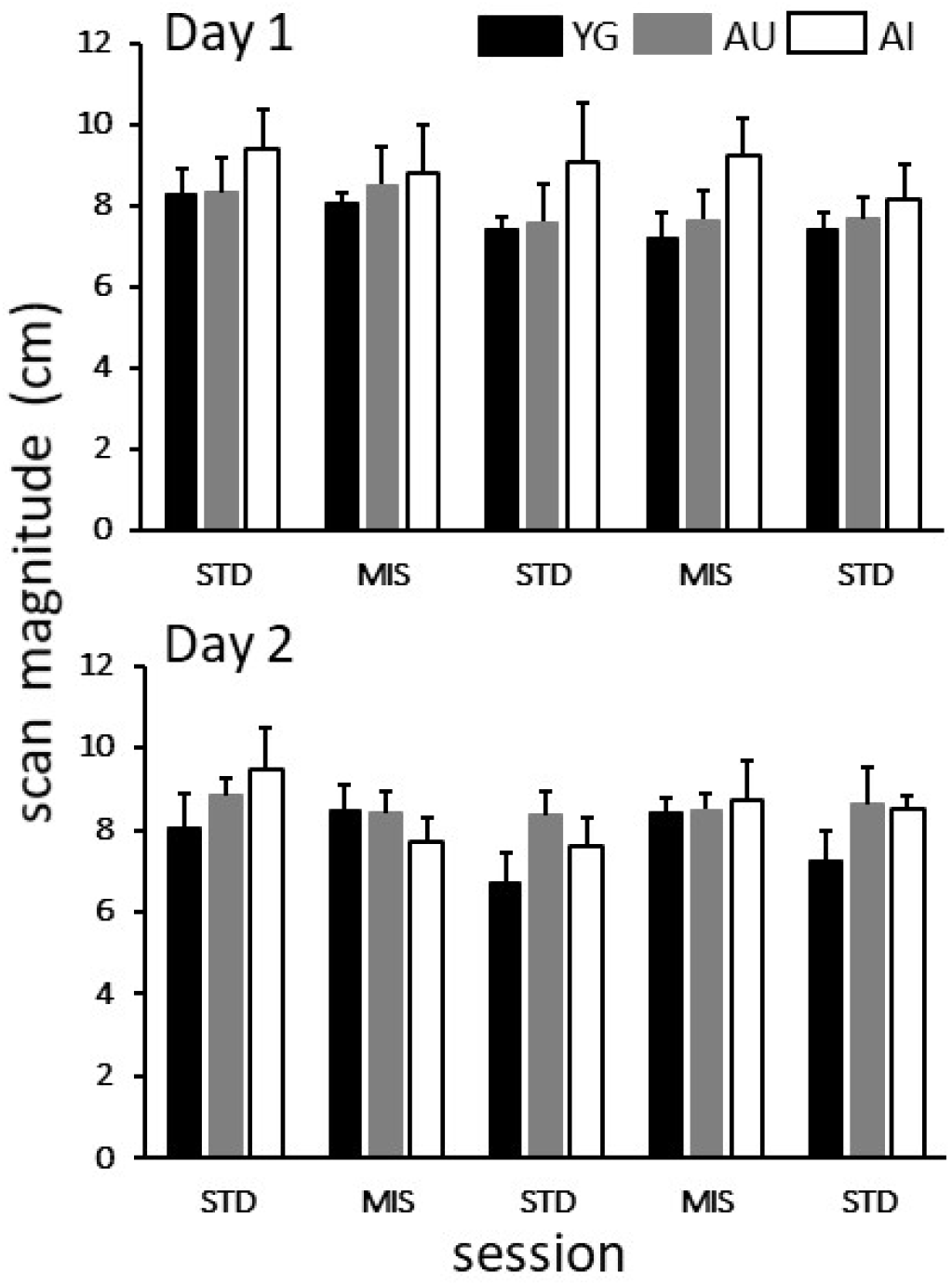
Scan magnitude across the 5 sessions (3 standard sessions interleaved with 2 mismatch sessions for Day 1 (top) and Day 2 (bottom) for YG (Black), AU (gray), and AI (white) animals. No significant main effects of age or session and no significant interaction were seen for Day 1. A significant main effect of session was shown for Day 2, but the interaction was not significant and none of the post hoc Tukey tests were significant (Supplementary Table 3).

**Supplementary Table 1.**
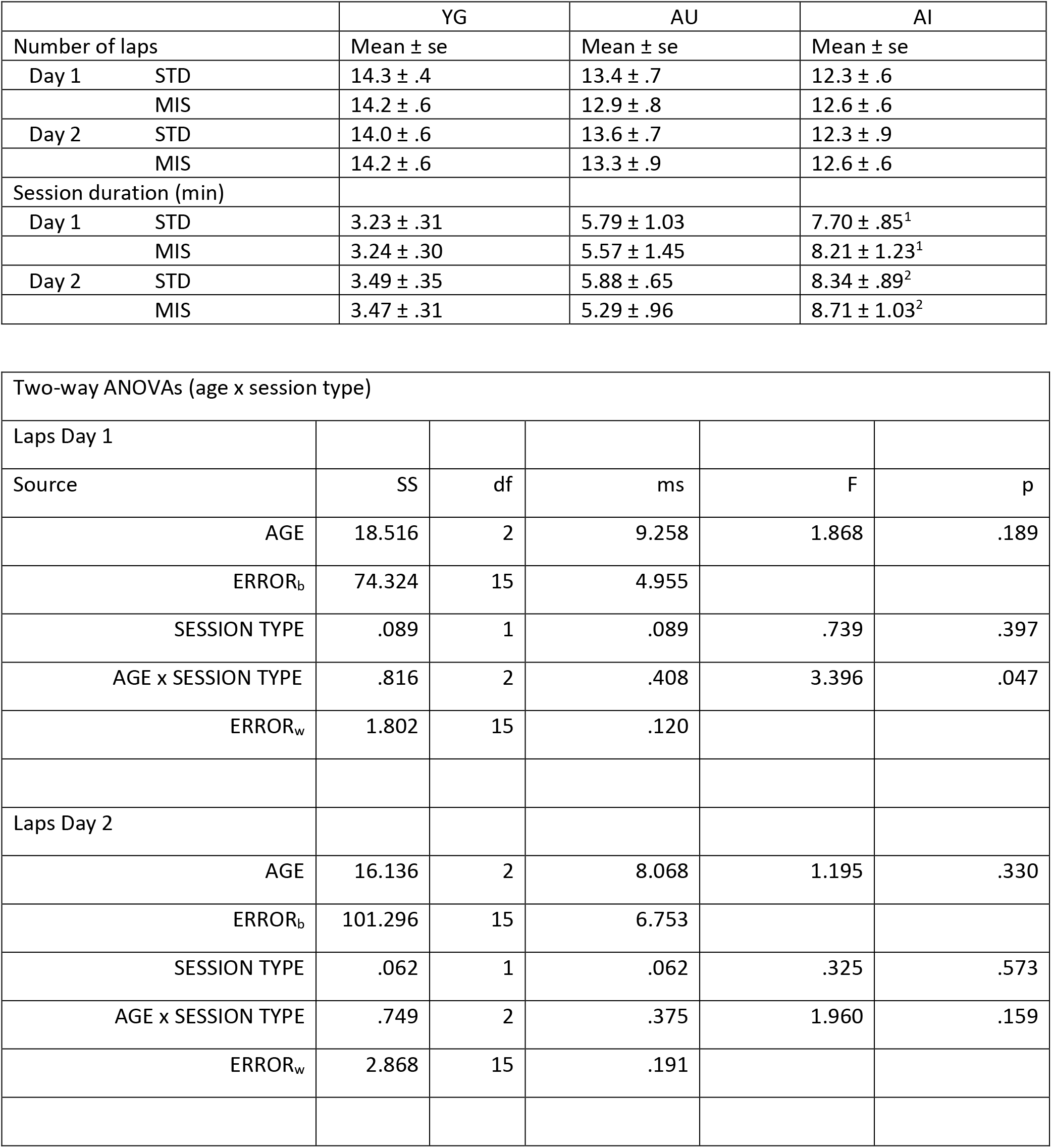
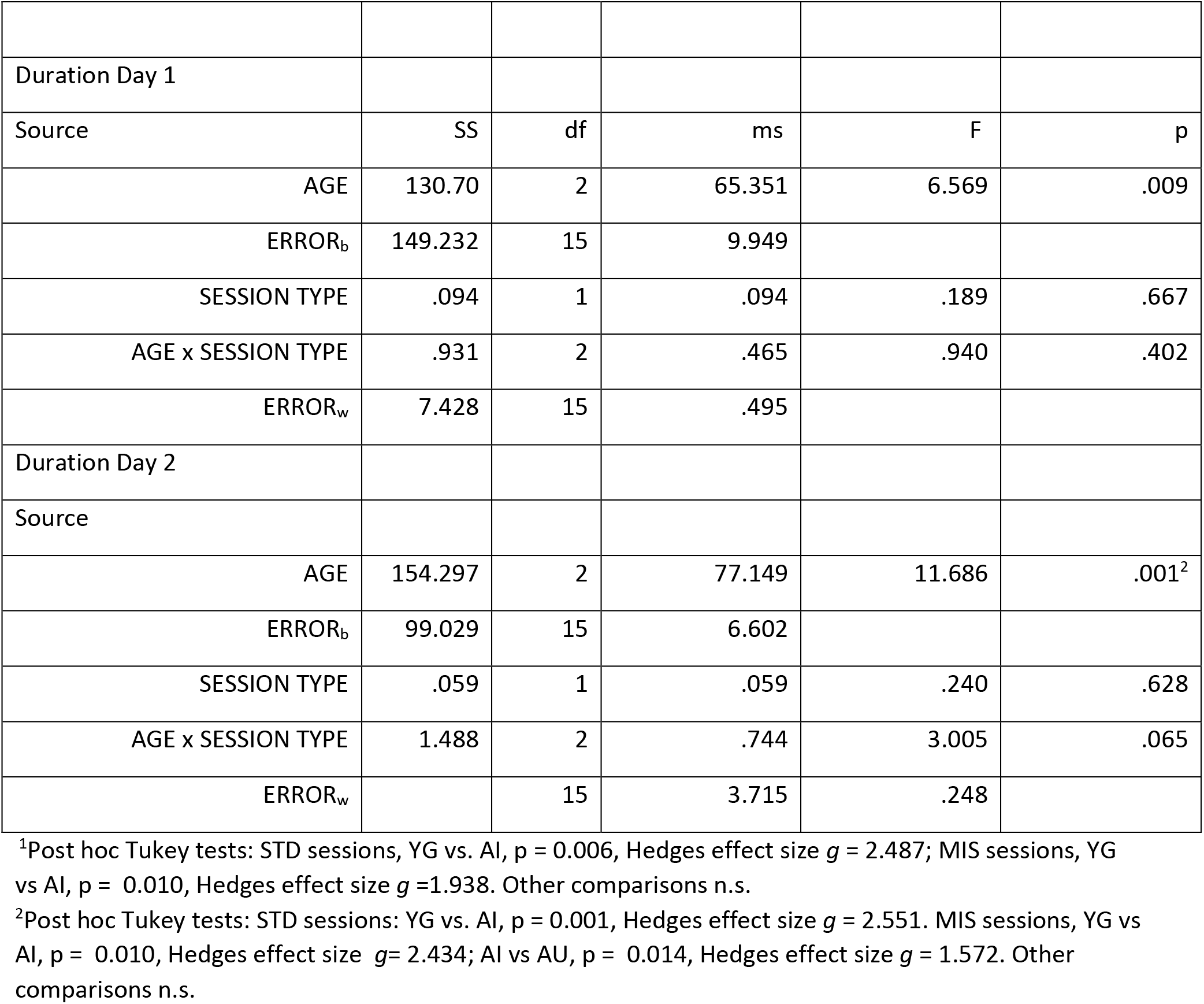
Effects of Age and Session Type (STD vs MIS) on number of laps and session duration. No significant effect of session type (mean of 3 STD sessions vs. mean of 2 MIS sessions) was observed upon number of laps or session duration. Session durations were significantly longer for the AI than for the YG rats regardless of session type on Day 1. On Day 2, AI STD session durations were significantly longer than YG rats, and AI MIS sessions were longer than YG and AU sessions.

**Supplementary Table 2.** Proportion of time spent scanning. There were no differences among age groups in baseline proportion of time spent scanning in Session 1. There was a significant effect of session number, as animals spent decreasing proportions of time scanning in later sessions of a day, but there were no effects of age and no significant age x session interactions.

|  | YG | AU | AI | ANOVA |  |
| --- | --- | --- | --- | --- | --- |
| Day 1, Session 1 only | 0.260 ± 0.037 | 0.296 ± 0.071 | 0.224 ± 0.030 | F(2,15) = 0.58, p = 0.57 |  |
| Day 2, Session 1 only | 0.259 ± 0.052 | 0.325 ± 0.074 | 0.241 ± 0.033 | F(2,15) = 0.69, p = 0.52 |  |
| Two-way ANOVAs (Age x Session) |  |  |  |  |  |
| DAY 1 |  |  |  |  |  |
| Source | SS | df | ms | F | p |
| AGE | 0.082 | 2 | 0.041 | 0.509 | 0.611 |
| ERROR<sub>b</sub> | 1.212 | 15 | 0.081 |  |  |
| SESSION | 0.032 | 4 | 0.008 | 3.101 | 0.020 |
| AGE x SESSION | 0.038 | 8 | 0.005 | 1.802 | 0.090 |
| ERROR<sub>w</sub> | 0.156 | 60 | 0.003 |  |  |
| DAY 2 |  |  |  |  |  |
| Source | SS | df | ms | F | p |
| AGE | 0.178 | 2 | 0.089 | 1.462 | 0.263 |
| ERROR<sub>b</sub> | 0.915 | 15 | 0.061 |  |  |
| SESSION | 0.040 | 4 | 0.010 | 2.472 | 0.052 |
| AGE x SESSION | 0.036 | 8 | 0.004 | 1.107 | 0.368 |
| ERROR<sub>w</sub> | 0.241 | 60 | 0.004 |  |  |

**Supplementary Table 3.** Scan magnitude ANOVA table

|  |  |  |  |  |  |  |
| --- | --- | --- | --- | --- | --- | --- |
| DAY 1 |  |  |  |  |  |  |
| Source |  | SS | df | ms | F | p |
|  | AGE | 27.082 | 2 | 13.541 | 0.704 | 0.510 |
|  | ERROR<sub>b</sub> | 288.526 | 15 | 19.235 |  |  |
|  | SESSION | 9.362 | 4 | 2.341 | 1.771 | 0.144 |
|  | AGE x SESSION | 6.113 | 8 | 0.764 | 0.578 | 0.793 |
|  | ERROR<sub>w</sub> | 79.288 | 60 | 1.321 |  |  |
| DAY 2 |  |  |  |  |  |  |
| Source |  | SS | df | ms | F | p |
|  | AGE | 8.482 | 2 | 4.241 | 0.463 | 0.638 |
|  | ERROR<sub>b</sub> | 137.341 | 15 | 9.156 |  |  |
|  | SESSION | 15.195 | 4 | 3.799 | 2.841 | 0.030 |
|  | AGE x SESSION | 13.334 | 8 | 1.667 | 1.246 | 0.285 |
|  | ERROR<sub>w</sub> | 80.237 | 60 | 1.337 |  |  |

**Supplementary Table 4.** The order in which the 4 possible cue mismatch angles (45°, 90°, 135°, 180°) were presented to each animal on Day 1 and Day 2 is shown in each row. The mean angle and s.e. for each age group (YG, AU, AI) and results of one-way ANOVAs within mismatch session are indicated below the data. No significant differences with age group were found within session in the randomly selected angles.

|  |  | Day 1 |  | Day 2 |  |
| --- | --- | --- | --- | --- | --- |
| rat | age | SESS 2 | SESS 4 | SESS 2 | SESS 4 |
| 1 | YG | 45 | 90 | 180 | 135 |
| 2 | YG | 45 | 180 | 135 | 90 |
| 3 | YG | 180 | 90 | 45 | 135 |
| 4 | YG | 180 | 45 | 135 | 90 |
| 5 | YG | 180 | 90 | 135 | 45 |
| 6 | AU | 90 | 45 | 135 | 180 |
| 7 | AU | 90 | 45 | 135 | 180 |
| 8 | AU | 135 | 45 | 180 | 90 |
| 9 | AU | 90 | 135 | 180 | 45 |
| 10 | AU | 90 | 180 | 45 | 135 |
| 11 | AU | 90 | 45 | 135 | 180 |
| 12 | AI | 135 | 90 | 45 | 180 |
| 13 | AI | 180 | 45 | 90 | 135 |
| 14 | AI | 135 | 90 | 45 | 180 |
| 15 | AI | 180 | 45 | 90 | 135 |
| 16 | AI | 180 | 135 | 45 | 90 |
| 17 | AI | 45 | 180 | 135 | 90 |
| 18 | AI | 135 | 90 | 180 | 45 |
|  |  | Day 1 |  | Day 2 |  |
|  | mean ± se | SESS 2 | SESS 4 | SESS 2 | SESS 4 |
|  | YG | 126.0 ± 33.1 | 99.0 ± 22.0 | 126.0 ± 22.0 | 99.0 ± 16.8 |
|  | AU | 97.5 ± 7.5 | 82.5 ± 22.4 | 135.0 ± 20.1 | 135.0 ± 23.2 |
|  | AI | 141.4 ± 18.1 | 96.4 ± 18.2 | 90.0 ± 19.6 | 122.4 ± 18.9 |
|  | age group ANOVA | F (2,15) = 1.266, p = 0.310 | F (2, 15) = 0.166, p = 0.848 | F (2,15) = 1.454, p = 0.265 | F (2, 15) = 0.729, p = 0.499 |

